# Rainbow-seq: combining cell lineage tracking with single-cell RNA sequencing in preimplantation embryos

**DOI:** 10.1101/293134

**Authors:** Fernando Biase, Qiuyang Wu, Riccardo Calandrelli, Marcelo Rivas-Astroza, Shuigeng Zhou, Sheng Zhong

**Author notes:** Equal contribution (Fernando Biase, Qiuyang Wu).

## Abstract

Single-cell RNA-seq experiments cannot record cell division history and therefore cannot directly connect intercellular differences at a later developmental stage to their progenitor cells. We developed Rainbow-seq to combine cell division lineage tracing with single-cell RNA-seq. With distinct fluorescent protein genes as lineage markers, Rainbow-seq enables each single-cell RNA-seq experiment to simultaneously read single-cell transcriptomes and decode the lineage marker genes. We traced the lineages deriving from each blastomere in two-cell mouse embryos and observed inequivalent contributions to the embryonic and abembryonic poles in 72% of the blastocysts evaluated. Rainbow-seq on four- and eight-cell embryos with lineage tracing triggered at two-cell stage exhibited remarkable transcriptome-wide differences between the two cell lineages at both stages, including genes involved in negative regulation of transcription and signaling. These data provide critical insights on cell fate choices in cleavage embryos. Rainbow-seq bridged a critical gap between cellular division history and single-cell RNA-seq assays.

## INTRODUCTION

A central question to developmental biology is how cells break molecular and functional symmetry during mitotic divisions. Two models have been proposed (Altschuler and Wu, 2010). In one model, a progenitor cell first divides into a set of homogeneous cells, and then these homogenous cells exhibit different tendencies towards different routes of differentiation (homogenous model). In the other model, every division creates a pair of cells with slight differences in their molecular profile, and accumulation of these differences lead to cell fate commitment (non-homogenous model).

Mammalian preimplantation development from a zygote to a blastocyst consisting of the functionally differentiated inner cell mass (ICM) and trophectoderm (TE) (Wolpert, 2015) is a perfect example of the symmetry-breaking paradigm. The first cell fate decision in mammals was believed to be an example of the homogenous model (Hiiragi et al., 2006). After fertilization, a mouse zygote was thought to undergo two or more rounds of symmetric cell divisions which create 4 or more homogeneous embryonic cells (blastomeres) (Leung and Zernicka-Goetz, 2015; Motosugi et al., 2005; Zernicka-Goetz et al., 2009). These homogeneous cells further divide and through a stochastic process (Wennekamp and Hiiragi, 2012) eventually self-organize into ICM and TE (Wennekamp et al., 2013) (Figure 1A). However, recent work on single-cell transcriptome profiling reported consistent gene expression differences between sister blastomeres in 2- and 4-cell mouse embryos (Biase et al., 2014), offering support to the non-homogeneous model (Hiiragi and Solter, 2006; Vogel, 2005).

**Figure 1.**
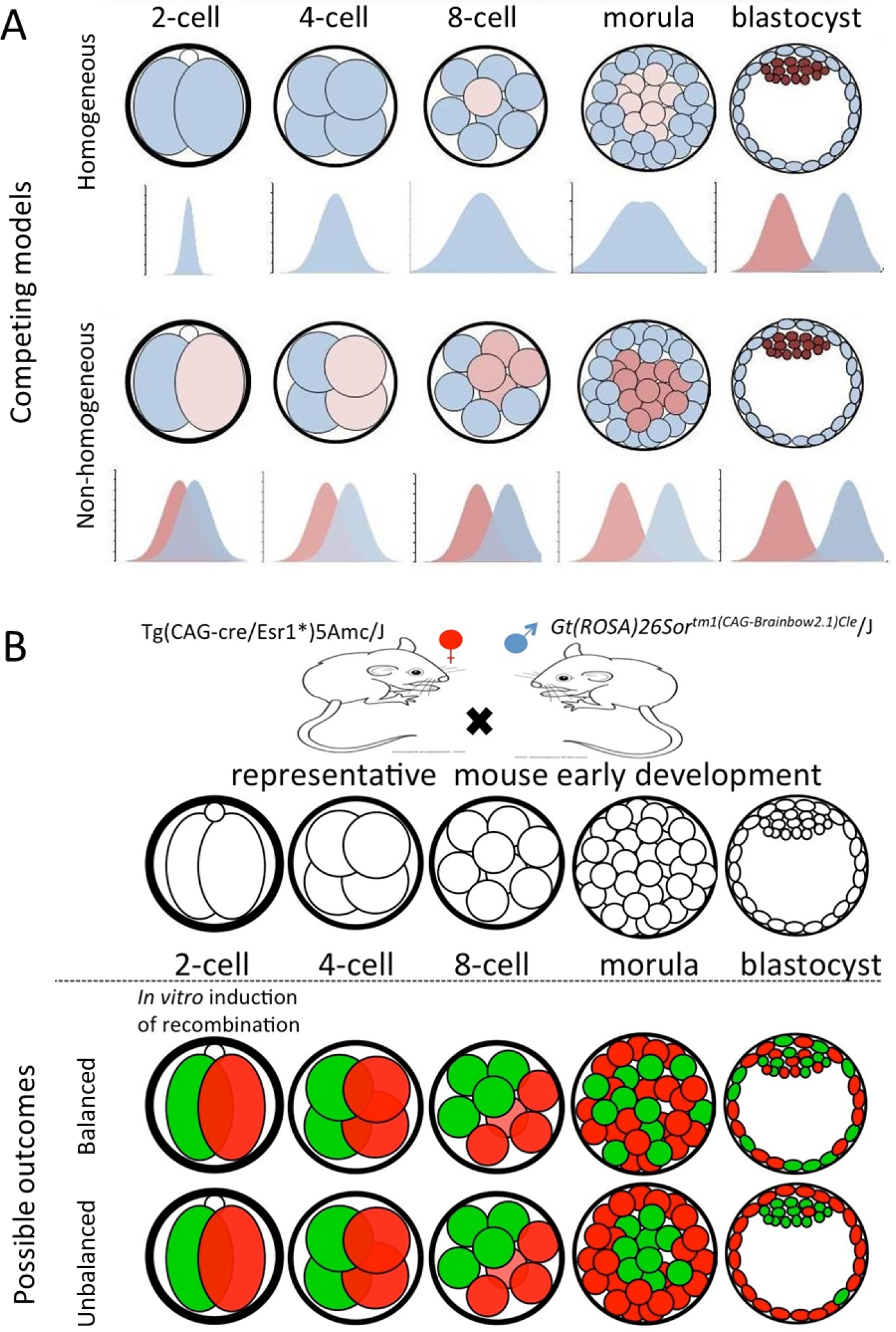
Competing models and lineage tracking strategy. (A) Homogeneous and non-homogeneous models differ in whether the 2 blastomeres at 2-cell stage have identical tendencies to contribute their descendent cells to inner cell mass (brown) and trophectoderm (blue-grey). (B) Cell lineage tracing was enabled by creation of preimplantation embryos heterozygous for brainbow sequence and Cre, and induction of Cre at 2-cell stage. At blastocyst stage, activated brainbow genes can exhibit either relatively equal (balanced) or biased (unbalanced) distributions in inner cell mass and trophectoderm.

The critical missing information for using single-cell transcriptome profiling to test the competing models is the history of cell divisions (Goolam et al., 2016; Plachta et al., 2011; Torres-Padilla et al., 2007). We approached the challenge of connecting cell-to-cell transcriptome differences at later embryonic stages to progenitor cells by developing Rainbow-seq, which combines cell division lineage tracking (Tabansky et al., 2013) with single-cell RNA sequencing.

## DESIGN

Advances in cell-specific genomic barcodes are revolutionizing cell lineage tracing. Leveraging genome editing tools (Kalhor et al., 2017; McKenna et al., 2016; Perli et al., 2016) or combinatorial DNA recombination loci (Polylox) (Pei et al., 2017), researchers were able to insert cell-specific exogenous DNA barcodes to the genomes of progenitor cells. The lineage information is propagated by DNA duplication and can be recovered in the progeny by targeted sequencing of the engineered genomic sequences. We reasoned that combining cell-specific genomic barcodes that express marker genes and single-cell RNA-seq would reveal single-cell transcriptome together with cell division lineage. To maximize the chances of obtaining expressible genomic barcodes, we resorted to the well tested Brainbow-2.1 construct that can express one of four fluorescent protein genes (GFP, CFP, RFP, YFP) (Livet et al., 2007). These fluorescent protein genes interspersed with recombination sites serve as both genomic barcodes and expressible cell markers. Their expression can be examined microscopically, offering an intermediate step of validation before single-cell RNA-seq analysis.

## RESULTS

### Cell lineage tracing starting with 2-cell embryos

We generated mouse embryos by crossing a strain that is homozygote for the brainbow sequence (Cai et al., 2013) with a mouse strain that is homozygote for a tamoxifen inducible Cre recombinase gene (Figure 1B). Tamoxifen treatment was expected to induce Cre translocation to the nucleus, which in turn could induce recombination on loxP sites, and thus lead to activation of one and only one of the four fluorescent protein genes in the brainbow sequence (Cai et al., 2013). We tested whether these anticipated effects could be achieved by a short application of tamoxifen to early 2-cell embryos (Figure S1A). Upon collection of early 2-cell embryos (42 h.p.i.) and a 2-hour 4-hydroxytamoxifen treatment, Cre protein translocated to the nuclei (Figure S1B), and within 1 hour post treatment the nuclear concentration of Cre reduced to background level (Figure S1B). Embryos undergone this brief tamoxifen treatment exhibited expression of fluorescent proteins at later developmental stages including 4-cell, 8-cell and blastocyst stages (Figure 2A). Hereafter, we name the hybrid embryos collected at early 2-cell stage and undergone a 2-hour 4-hydroxytamoxifen treatment as *rainbow embryos*, and call the Cre-activated gene as *lineage marker gene*.

**Figure 2.**
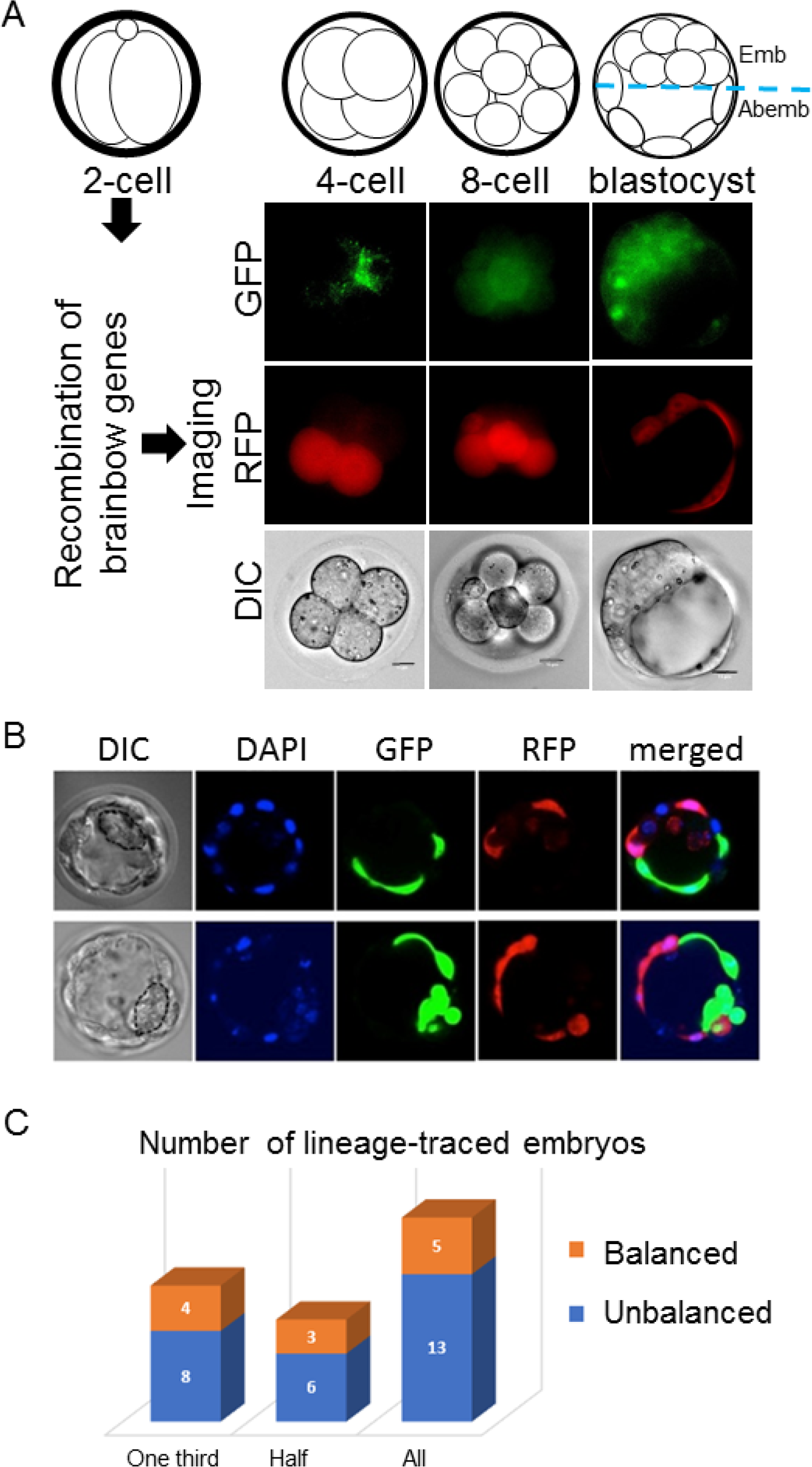
Cell lineage tracking in mouse preimplantation embryos. (A) Recombination of brainbow construct was induced in early 2-cell embryos, and images were acquired at 4-, 8-cells and blastocyst stages. GFP: green fluorescent protein; RFP: red fluorescent protein; DIC: differential interference contrast. Scale bar: 15um. Image was acquired by a wild-field microscope containing filters for RFP and GFP. (B) Two representative blastocyst stage embryos imaged with DIC, DAPI, GFP, and RFP channels. (C) Numbers of blastocyst-stage embryos with balanced (orange) and unbalanced (blue) lineage marker expression in the two embryonic poles, with respective counting methods (three lanes). Data from different counting methods are shown in columns, which include requiring at least 1/3 cells (One third) or 1/2 cells (One half) expressing either GFP or RFP, and all the imaged embryos (All).

### Daughter cells of blastomeres at 2-cell stage tend to contribute inequivalently to embryonic and abembryonic poles at the blastocyst stage

We leveraged the brainbow embryos to revisit the question on whether the daughter cells produced by each blastomere of a 2-cell embryo contribute equivalently to embryonic and abembryonic poles in blastocyst-stage embryos (Hiiragi and Solter, 2004; Plusa et al., 2005; Vogel, 2005). Our experiment was designed as follows. First, a total of N blastocyst-stage rainbow embryos would be imaged by a wide-field fluorescent microscope with filters for GFP, RFP, and DAPI (Figure 2A-B). Because we had to devote one of three channels to nuclei staining (DAPI) for cell counting, this experimental design would not be able to detect CFP and YFP expressing cells. Next, the imaged embryos that had more than 1/3 of their cells in the embryonic pole expressing a detectable lineage marker (either GFP or RFP) would be used for data analysis. A contingency table would be built for each embryo documenting the numbers of GFP expressing (GFP+) and non-GFP expressing (GFP−) cells in the embryonic pole and in the abembryonic pole. An odds ratio (OR) would be calculated for each embryo, and if 1/3 < OR < 3, this embryo is regarded as having balanced GFP+ cells in the two poles, otherwise the embryo is regarded as having unbalanced GFP+ cells in the two poles. The same analysis would be carried out for RFP.

We imaged 18 early blastocysts expressing RFP and/or GFP, of which 12 embryos had more than 1/3 of cells expressing fluorescent protein. Eight of those embryos (66%) exhibited unbalanced GFP+ or RFP+ cells in the two poles (first column, Figure 2C). For sensitivity analyses, we used two alternative counting methods to re-estimate the proportion embryos with unbalanced distribution of a cell lineage between the two poles. First, we required the embryos to be included for analysis to have at least 50% of its cells expressing either GFP or RFP, resulting in a total of 9 embryos, of which 6 embryos (66%) exhibited unbalanced GFP+ or RFP+ cells in the two poles (second column, Figure 2C). Second, we counted in GFP+ cells in the embryonic pole (n1) and GFP+ cells in the abembryonic pole (n2) and regarded any embryo with 1/3 < n1/n2 < 3 as having balanced GFP+ cells, and did the same counting for RFP. All 18 embryos were included in this analysis, of which 13 embryos (72%) were unbalanced (third column, Figure 2C). Taken together, approximately two thirds rainbow embryos exhibited unbalanced distribution of the two cell lineages in the two embryonic poles. Previous efforts obtained discrepant estimates of such a proportion (Hiiragi and Solter, 2004; Piotrowska et al., 2001; Plusa et al., 2005; Vogel, 2005). Our results obtained from rainbow embryos were better aligned with that suggesting more embryos exhibiting unequal contribution of the two lineages than those exhibiting equal contribution to the embryonic and abembryonic poles.

### Single-cell RNA sequencing analysis of rainbow embryos

We induced lineage tracking from 2-cell stage and carried out Rainbow-seq on nine 4-cell embryos and four 8-cell embryos. We lost one blastomere from an 8-cell embryo during the manipulation, and produced paired-end 100nt sequencing data from a total of 67 single blastomeres, including 36 from 4-cell stage and 31 from 8-cell stage (Tables S1, S2). One 8-cell stage blastomere did not pass our quality control due to producing fewer than 3 million uniquely mapped sequencing reads (Row E13, B7, Table S2). The remaining 66 cells yielded on average 21 million uniquely mapped read pairs (mm10) per cell (Figure S2A). We used single-cell RNA-seq reads mapped to the brainbow 2.1 sequence to determine which lineage marker gene was expressed, and categorized the cells from each embryo into two groups (lineage A and B) based on its expressed marker gene (Lineage column, Tables S1, S2). Data was insufficient to resolve three 4-cell stage embryos (12 blastomeres) and four 8-cell stage blastomeres due to limited sequencing reads mapped onto any lineage marker gene. We deposited all sequencing data into GEO (GSE106287) for public access, however we limited our analyzes to the 50 blastomeres whose lineage can be traced to either one of the sister blastomeres at 2-cell stage.

The first two principle components explained 8% of sample variations, where the developmental stage was a major source of variation (Figure S2B). With FPKM>1 as the threshold for calling expressed genes, the 4-cell stage blastomeres on average expressed 6,852 coding genes per cell. A total of 15,218 coding genes were expressed in a least one 4-cell stage blastomere. The 8-cell blastomeres on average expressed 6,474 coding genes per cell, with a total of 15,342 coding genes expressed in at least one 8-cell blastomere.

### Transcriptome-wide expression difference between cell division lineages

Recognizing that data of individual genes may not provide conclusive evidence to either the homogenous or the non-homologous model, we analyzed data of the entire transcriptome. We carried out two transcriptome-wide tests. First, we tested whether there is non-zero probability for any subset of genes to exhibit greater between-lineage variation than embryo-to-embryo variation. To this end, we fit a generalized linear model to every gene (*g*), and calculated the sum of squares to account for the between-lineage variation 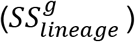 and embryo-to-embryo variation 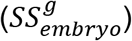. For every gene, we calculated the ratio (*r*^*g*^) of lineage sum of squares to combined sum of squares of embryo and lineage. The empirical distribution of this ratio (denoted as *R*) exhibited greater tail probabilities than a series of three β distributions, β(2,5), β(1,3), and β(5,5), each of which has non-zero probability in any non-singular interval near 1 ([1−δ,1], (δ>0)) (Figure 3A-D). The non-zero probability for a subset of genes to exhibit greater lineage variation than embryo variation suggests a transcriptome-wide difference between the two division lineages.

**Figure 3.**
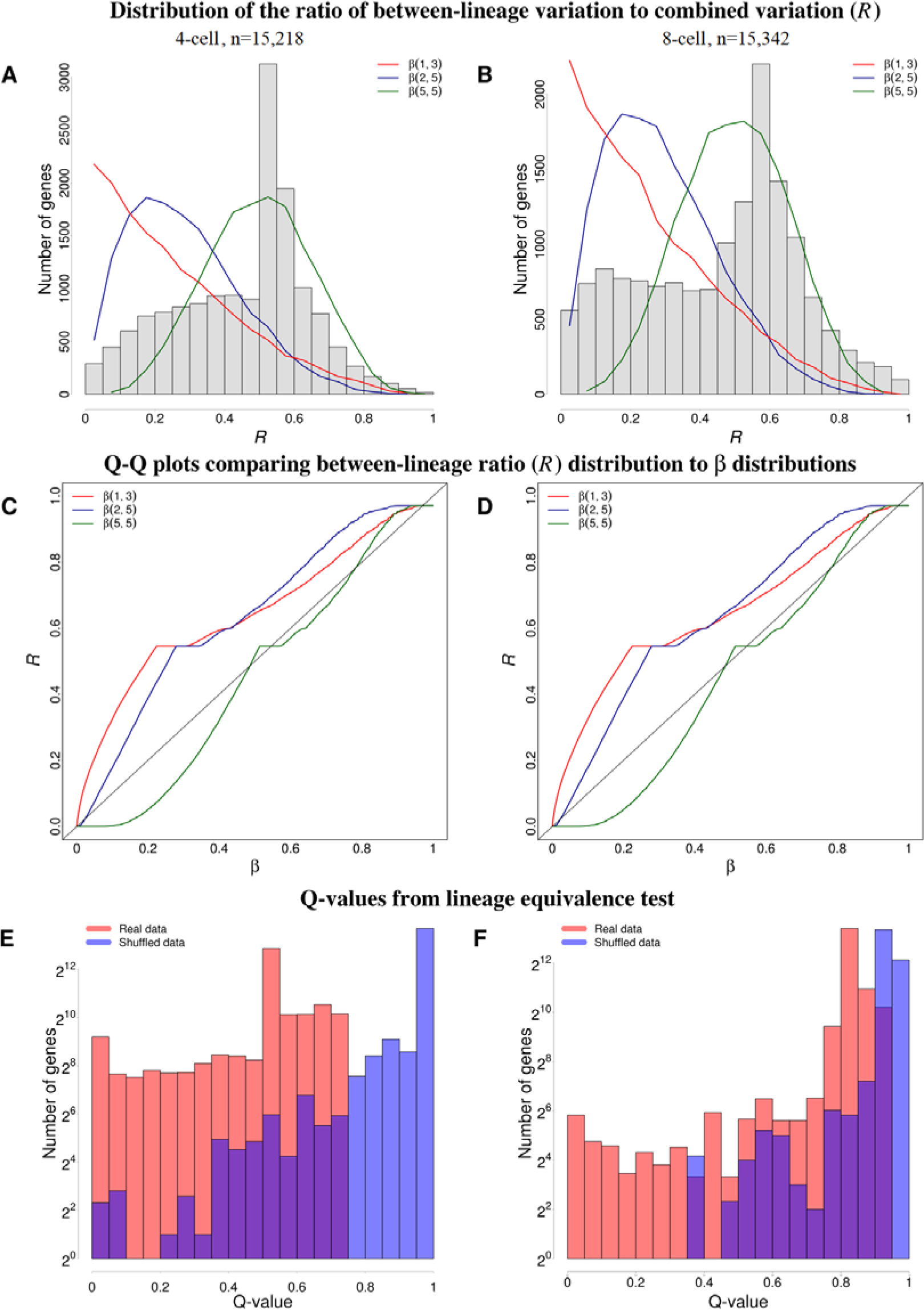
Transcriptome-wide differences between two lineages. (A-B) Empirical distributions of random variable *R* (ratio of between-lineage variation to combined variation) in 4-cell (A) and 8cell stage embryos (B), superimposed with a series of β distributions (red: β(1,3), blue: β (2,2), green: β(5,5)). (C-D) Q-Q plots of *R* versus a series of β distributions in 4-cell (C) and 8-cell stage embryos (D), where red, blue, and green represents β(1,3), β (2,2), and β(5,5), respectively. (E-F) Distributions of q-values from lineage equivalence tests based from real data (red) and shuffled data (blue) in 4-cell (E) and 8-cell stage blastomeres (F).

Second, after a gene-by-gene test of the null hypothesis that this gene does not exhibit between-lineage expression differences (ANOVA, Methods), we obtained the empirical distribution of the q-values for all the genes from such a test (red bars, Figure 3E-F). To derive a background distribution, we shuffled the lineage labels on every blastomere and carried out the same test (blue bars, Figure 2E-F). The distribution of q-values from real data was skewed towards lower values as compared to the q-values from shuffled data (p-value < 2.2e-16, Kolmogorov test), reflecting a transcriptome-wide difference between the two cell division lineages.

### Specific genes with expression differences between cell division lineages

The genes with the largest between-lineage differences in transcript abundance at 4-cell stage were involved in basic transcription machinery including *Gtf2f1* (General Transcription Factor IIF Subunit 1, TFIIF) and *Eloa* (RNA Polymerase II Transcription Factor SIII Subunit A1, SIII), negative regulations of P53 activity (*Aurkb*) and WNT signaling (*Ctnnbip1*, Inhibitor of Beta-Catenin and Tcf4), nuclear pore complex (*Nup35*), protein synthesis in cytoplasm (*Rbm3*) and in mitochondrion (*Mrpl10*), endoplasmic reticulum biogenesis (*Atl2*), and glycogen synthesis (*Fbp2*) (Figure 4A). The genes with the largest between lineage variations at 8-cell stage were transcriptional suppressors *Bclaf1* and *Smad7*, the chromatin remodeler *Atrx* involved in deposition of H3K9me3, histone demethylase *Kdm2b* related to removal of active histone marks including H3K4me3, vesicle trafficking and secretory pathways including Golgi membrane proteins *Glg1* and *B4galnt2*, trans-Golgi network *Snx15* and Golgi-to-ER transport *Stx8* (vesicle fusion, Golgi-to-ER transport), cell shape modulator *Fez2*, and the lincRNA genes *Gm12514* and *GM14617* (Figure 4B). Interestingly, suppression of H3K4 demethylase *Kdm2b* in mouse oocytes resulted in impaired blastocyst formation (see extended Figure 6e in (Liu et al., 2016)). Even though the above-mentioned genes appeared to support the non-homologous model, it is our opinion that statistical tests based on the entire transcriptome could better assess the two competing models than that based on each individual gene.

**Figure 4.**
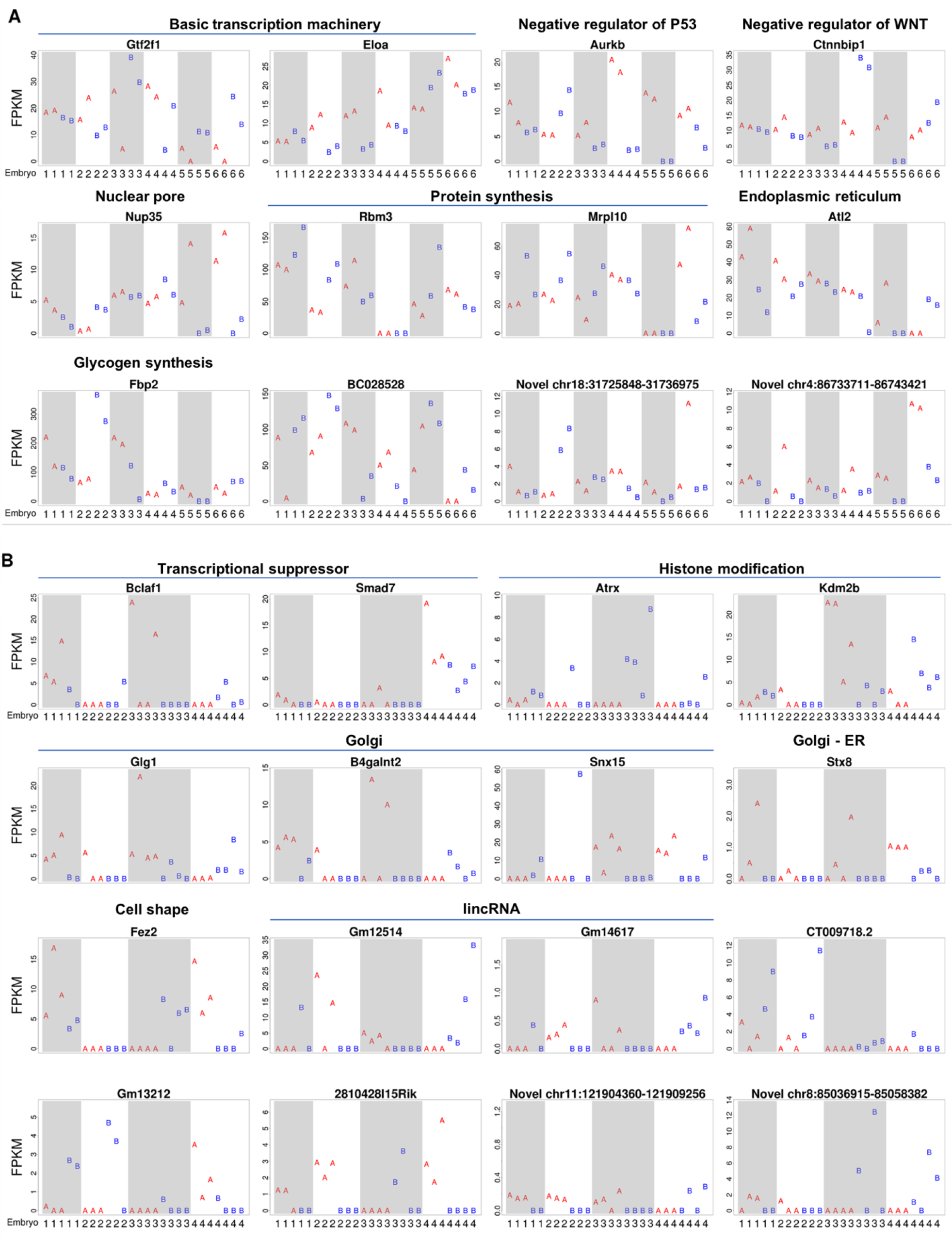
Genes with the between-lineage differences at 4- (A) and 8-cell stages (B). FPKM (y axis) of every blastomere (column) in each embryo (marked by embryo number in columns). Shaded columns delineate different embryos. The blastomeres of the two lineages are marked with A (red) and B (blue), respectively.

### Transposon related novel transcript isoforms

We tested whether any transcript isoforms were specifically expressed in early stage preimplantation embryos. To this end, we combined all the Rainbow-seq datasets to obtain over 2.1 billion paired-end reads, of which over 1.6 billion read pairs could be uniquely mapped to the genome (mm10). Using Cufflinks (v2.2.1 with -g parameter), we recorded a total of 20,371 novel transcript isoforms, corresponding to 7,594 annotated genes. Among these novel isoforms, 3,632 (17.8% of all novel isoforms) contained transposon sequences according to Repeatmasker database (February 2015 release) (Figure 5A), which will be referred to as <u>T</u>ransposon <u>Re</u>lated <u>N</u>ovel <u>I</u>soforms (TRENI). These 3,632 TRENIs corresponded to 1,701 Refseq genes, in which “poly(A) RNA binding”, “RNA binding”, and “nucleotide binding”, “nucleic acid binding” were enriched (Benjamini adjusted p-value < 1.1e-7) Gene Ontology functions (DAVID v6.8) (Figure 5B), suggesting a recurring theme in RNA binding and processing.

**Figure 5.**
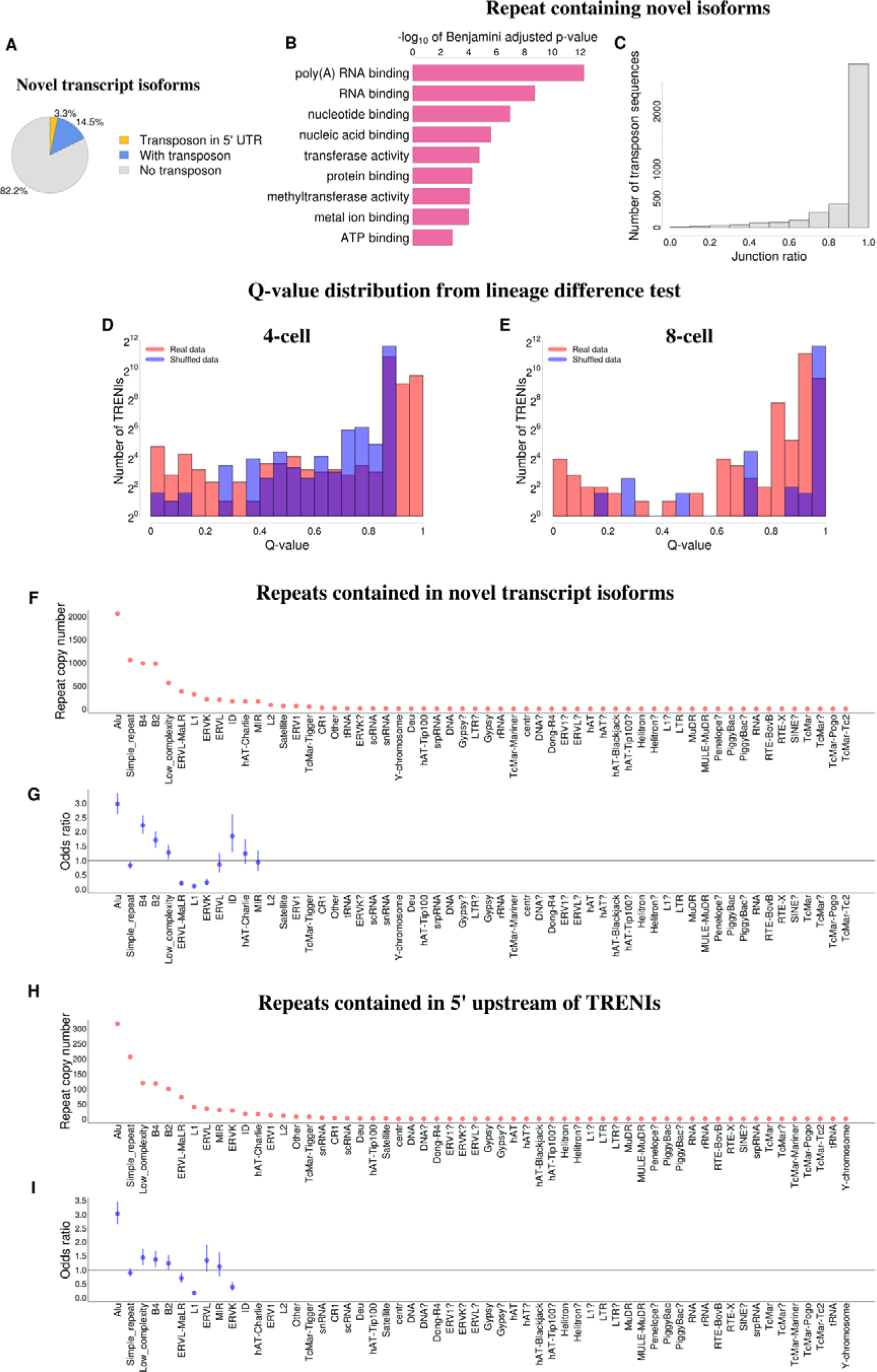
Transposon related novel isoforms (TRENIs). (A) Pie-chart of novel transcript isoforms, including 16,739 that do not contain any repeat sequence (grey), 3064 and 673 isoforms with transposons located downstream to start codons (blue) and in 5’ UTRs (yellow), respectively. (B) Enriched GO molecular functions in TRENIs, ranked by multiple hypothesis testing adjusted p-values. (C) Histogram of junction ratios for all TRENI contained transposons. (D-E) Histogram of TRENIs’ q-values derived from tests of equivalent expression between lineages at 4- (D) and 8-cell (E) stages, based from real data (red) and shuffled data (blue). (F) Copy number of repeats contained in TRENIs, categorized by transposon family (columns). (G) Odds ratios between TRENIs and each repeat family (columns). Vertical bars: 95% confidence intervals. Odds ratio was not calculated for those repeat families when the TRENIs associated with a repeat family were fewer than 20. (H) Copy number of repeats contained in TRENIs’ 5 UTRs, categorized by transposon family (columns). (I) Odds ratios between 5’UTR-TRENIs and each repeat family (columns). Vertical bars: 95% confidence intervals. Odds ratio was not calculated for those repeat families when the 5’UTR-TRENIs associated with a repeat family were fewer than 20.

We asked whether the incorporation of these transposons in TRENIs were supported by junction reads spanning across the transposon and the nearest known Refseq exons. To this end, we computed the ratio of junction reads and the total number of reads aligned to each transposon, hereafter denoted as junction ratio (JR, Methods). We identified 3,902 transposons contained in TRENIs, of which 2,817 (72.19%) exhibited JR of 0.9 or above (Figure 5C).

To test whether TRENIs exhibit betweenlineage expression difference, we derived a q-value for every of the 3,632 TRENIs by ANOVA (Methods). For comparison, we shuffled the lineage labels on all blastomeres and derived q-values again (background q-values). The distribution of real data q-values differed with that of background q-values (p-value < 2.2e-16, Kolmogorov test) (Figure 5D-E, Figure S3). More importantly, a total of 37 TRINEs exhibited q-values smaller than 0.01, whereas none of the background q-values were smaller than the same threshold (Figure S3). Finally, all the repeats expressed in 4- and 8-cell embryos, regardless of whether they were contained within TRINEs, did not exhibit statistically discernable between-lineage differences (the distribution of q-values derived from real data was not different from that of shuffled data) (Figure S4).

### Murine Alu makes the largest contribution to novel transcript isoforms

We asked whether different repeat families contributed equally to TRENIs. In the total of 58 repeat families (RepeatMasker) 18 families had 20 or more genomic copies included in TRENIs (Figure 5F). To account for genome-wide copy numbers of each repeat families, we calculated the odds ratio for each repeat family to be contained in TRENIs. When repeat families were ranked by this odds ratio, Alu, B4, ID, and B2 were most enriched in TRENIs (Figure 5G). We asked whether the association between any repeat family and TRENIs would became more pronounced when only high-confidence transposon-containing TRENIs were included in the analysis. To this end, we repeated the above analysis using only the 2,817 transposons with JR ≥ 0.9, which resulted in similar odds ratios with Alu being the most enriched repeat family (Figure S5).

Next, we asked whether different repeat families contributed equally to TRENIs’ 5’ UTR regions. A total of 673 (18.53%) TRENIs in which the transposons were inserted in the 5’ upstream to previously annotated start codons (5’UTR-TRENIs) (Figure 5H), among which 418 contained murine Alu, simple_repeat, low_complexity, B4, and B2 repeats in 5’ UTRs, where murine Alu alone accounted for more than 300 5’ UTRs (Figure 4H). We calculated odds ratio to quantify the association of transposons in each transposon family to 5’UTR-TRENIs. Murine Alu exhibited the largest enrichment to 5’ UTRs in these novel transcript isoforms (odds ratio =3.03, p-value =0, Chi-squared test) (Figure 4H). In contrast, the other repeat families exhibited either moderate enrichment (1 < odds ratio < 1.5) or depletion (odds ratio < 1) in 5’UTR-TRENIs (Figure 5I). Finally, 9 and 26 5’UTR-TRENIs exhibited expression differences between the two division lineages at 4- and 8-cell stages, respectively, including novel transcript isoforms of *Gata4* (Figure S6A) and *Khdc1a* (Figure S6B).

## DISCUSSION

### Lack-of-continuity Dilemma

Not a single gene has been reported to exhibit continuous between-lineage expression difference starting from the two blastomeres at 2-cell stage until the formation of ICM and TE at blastocyst stage. This is not completely unexpected considering that comprehensive transcriptome analyses revealed highly dynamic gene transcription in early stage mouse preimplantation embryos (Kidder and Pedersen, 1982; Xie et al., 2010), likely attributable to combined effects of cleavage-related split of RNA pool, RNA degradation and zygotic genome activation (Biase et al., 2014). The lack of identified single gene(s) with continuous expression divergence posed a dilemma to the non-homologous hypothesis, in that if the two blastomeres at 2-cell stage are poised for cell fate decisions, what could to be the molecular means to record and later exhibit such a difference? Hereafter we call this dilemma Lack-of-continuity Dilemma.

### Reconciliations to Lack-of-continuity Dilemma

The dilemma was rooted in the non-homogenous model and may be resolved in several ways. First, the cell-fate decision is initiated after the first cleavage. This idea essentially refutes the non-homogeneous model for the first cleavage. Second, the lineage difference is passed from one gene activated at an earlier stage, for example 2-cell stage, onto another gene activated at a later stage. Given the transcription rate and mRNA half-life are both an order of magnitude smaller than the time of a cleavage, this model is within the boundaries of biophysics principles. Third, the lineage difference is recorded in the order of expression levels of a pair or list of genes, and this order is preserved throughout preimplantation development. For example, at 2-cell stage the expression level order is A>B in one blastomere and B>A in the other blastomere, and such an order is preserved in inner cell mass and trophectoderm. The third explanation was supported by single-cell RNA-seq data (Biase et al., 2014). Taken together, there are plausible means to explain the dilemma while withholding the homologous model.

### Rainbow-seq approach and inference

Rainbow-seq addresses the Lack-of-continuity Dilemma from another perspective. The main idea is to combine cell division tracing with single-cell RNA-seq. A genetic cell-lineage tracer was activated at 2-cell stage and analyzed at later stages. This method has two advantages. First, activation of distinct tracer genes in each blastomere’s genome was achieved by a relatively simple procedure, thus removing the necessity of highly invasive procedure to microinject dyes into live embryos. Second, through a genetic tracer embedded in the genome, DNA duplication ensured inheritance of the tracer during cell divisions. We used Rainbow-seq to test the consequential events of two cell division lineages created by the first cleavage, and detected between-lineage difference in both 4- and 8-cell stages of mouse preimplantation embryos. The difference was most pronounced in the collective distribution of the transcriptome (Figure 3).

The genes with between-lineage differences appeared to emphasize a theme of negative regulation, including inhibitors of the WNT pathway *Ctnnbip1* (*Inhibitor of Beta-Catenin and Tcf4*) (Tago et al., 2000) and *Smad7* (Han et al., 2006); *Aurora Kinase B* that negatively regulates P53’s transcriptional activity by promoting its degradation (Gully et al., 2012); chromatin remodeler *Atrx* that promotes repressive histone mark H3K9me3 and formation of heterochromatin (Sadic et al., 2015); histone demethylase *Kdm2b* that removes active histone mark H3K4me3 (Tsukada et al., 2006); and glycosyltransferase *B4galnt2* involved in negative regulation of cell-cell adhesion (Kawamura et al., 2005) (Figure 4). Considering that negative feedback as exemplified in Delta-Notch signaling is perhaps the best known molecular mechanism for neighboring cells to take on divergent functions (Artavanis-Tsakonas et al., 1999), the emergence of negative regulators from Rainbow-seq analysis may allude to cell-cell communication between the first two cell lineages. The other emerged theme of vesicle trafficking and secretion as exemplified by *Atl2, Glg1, B4galnt2, Snx15*, and *Stx8* is consistent with the idea of cell-cell communication at very early embryonic stages (Calarco-Gillam, 1986) (Sozen et al., 2014). Finally, *Fbp2’s* divergent presence in the two lineages is reminiscent to a sperm-related embryo patterning hypothesis (Takaoka and Hamada, 2014; Zernicka-Goetz, 2002), considering presence of *Fbp2* mRNA in mouse sperm (Schuster et al., 2016) and interactions of FBP2 and ALDOA, a sperm-head protein essential for sperm-egg interaction (Petit et al., 2013).

The novel transcript isoforms of known genes that contained repeat sequences (TRENIs) exhibited lineage difference, especially at the 8-cell stage (Figure 5D). At 8-cell stage, chromatin remodeler *Atrx* that responds to DNA demethylation and suppresses expression of hypomethylated repeat sequence in preimplantation embryos (He et al., 2015) exhibited between lineage expression difference (Figure 4B). These data suggest a model where DNA demethylation on repeat sequence is asymmetrically compensated by ATRX in the two cell lineages, leading to differential expression of transposon-containing transcripts (Figure 6). Alu repeats were most enriched in TRENIs, and the murine Alu family was the only repeat family with significant enrichment to 5’ UTR of TRENIs. These expression data may offer an explanation to the selective retention of Alu repeats to upstream and intronic regions of known genes (Tsirigos and Rigoutsos, 2009), where the selectively retained Alu repeats are part of rare transcript isoforms, which are specifically expressed in preimplantation embryos. Furthermore, asymmetric suppression of these repeats leads to asymmetric expression of genes required for the first cell fate decision, which is the functional force for selective retention. Consistent to this idea, the same group of genes involved in “RNA binding” were enriched in TRENIs and were reported by an independent analysis as the neighboring genes to selectively retained Alu repeats (Tsirigos and Rigoutsos, 2009).

**Figure 6.**
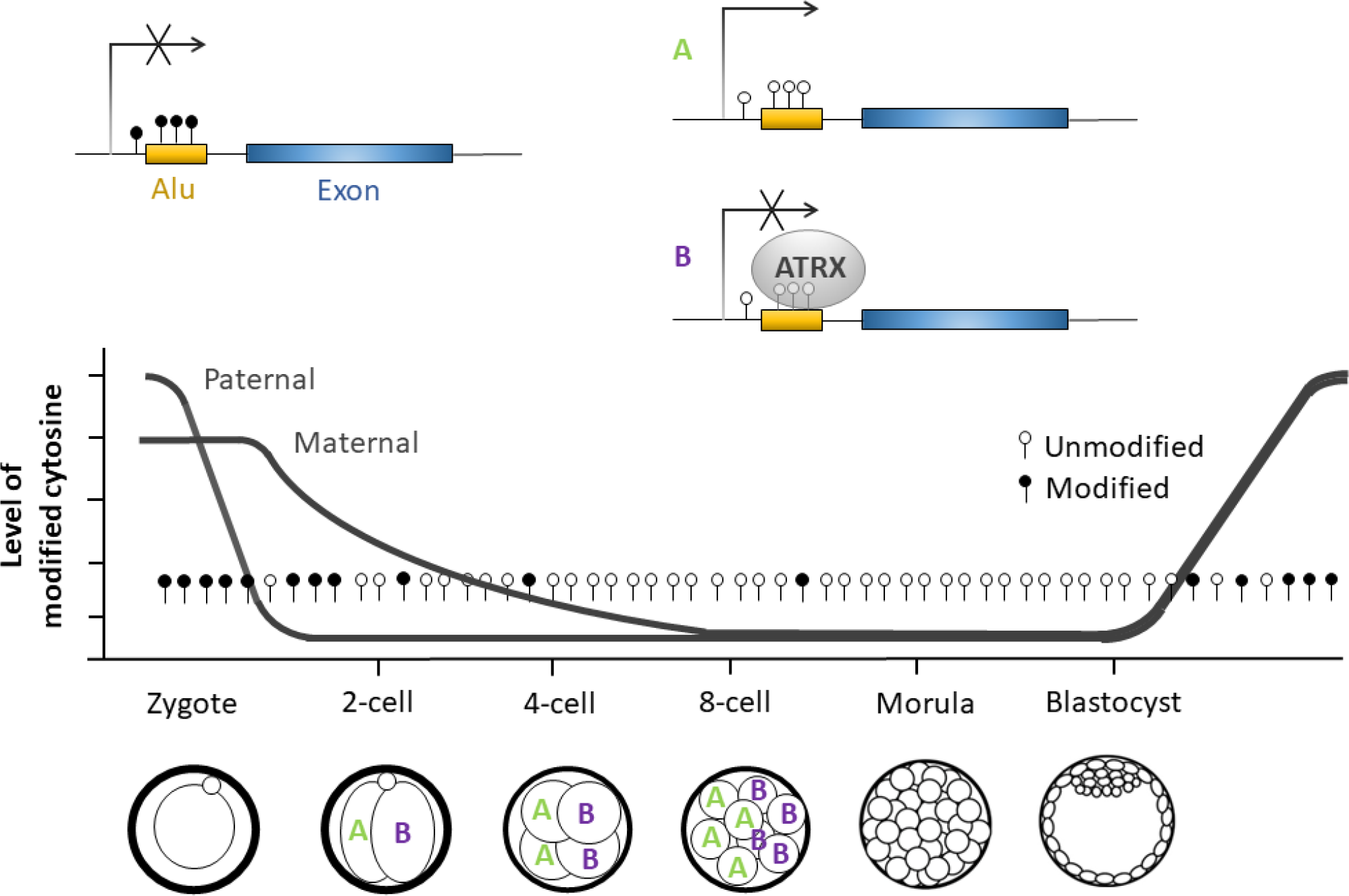
A model of repeat sequence related lineage difference. Decrease of modified cytosine (filled circles) allows repeat sequence (yellow) to be expressed (A). *Atrx* is expressed in one lineage where ATRX preferentially attaches hypomethylated (empty circles) repeats and suppresses their expression (B).

### Limitations

The number of fluorescent protein genes limits the number of lineages that can be resolved. This can be overcome by substituting fluorescent protein genes with random blocks of sequences (Pei et al., 2017), if the patterning of cell distribution is not relevant for the investigation prior to single-cell transcriptome analysis.

## MATERIAL AND METHODS

### Generation of rainbow mouse embryos

All the procedures for animal handling and breeding were approved by Institutional Animal Care and Use Committees of University of California San Diego. For the generation of rainbow embryos, we crossed two transgenic strains obtained from Jackson Laboratory. The strain STOCK Gt(ROSA)26Sortm1(CAG-Brainbow2.1)Cle/J (stock number 013731, RRID:SCR_004894) is homozygous for one copy of the brainbow construct v2.1, and the strain STOCK Tg(CAG-cre/Esr1*)5Amc/J (IMSR Cat# JAX:004453, RRID:IMSR_JAX:004453) is homozygous with one copy of the gene coding CRE protein. We crossed mice from each of the strains (1:1) to generate hybrid embryos containing one copy of the brainbow construct v2.1 and one copy of the Cre gene.

Adult female mice were estrous-synchronized and super-ovulated by two consecutive intra-peritoneal injections of hormones (to 2.5 IU of hCG and 5IU of PMSG, PG600, Intervet). The injections were administered 48 hours apart, and following the second injection, females and males were mated (1:1). Two-cell embryos were collected 46 hours post second hormonal injection (hpi) and incubated (37°C, 5%CO_2_, 100% humidity) for two hours with 4-Hydroxytamoxifen (100μm, Sigma) in KSOM media (Millipore).

### Embryo imaging

Tamoxifen-treated embryos were cultured *in vitro* (37°C, 5%CO_2_, 100% humidity) until reaching 4-cell, 8-cell or blastocyst stages (~ 54, 70 and 90hpi, respectively), briefly treated with NucBlue™ Live Cell Stain (Invitrogen) in KSOM media and taken to wild-field inverted fluorescent microscope (Olympus). Embryos were imaged in KSOM media, with filter cubes for DAPI (Ex: 350/50x, Em: 460/50m), GFP (Ex: 480/40x, Em: 535/50m), RFP (Ex: 545/25x, Em: 605/70m) and differential interference contrast, and a 40x dry objective (NA=0.95, WD=0.18mm). Images were captured with an ORCA R2 monochromatic CCD camera (Hamammatsu) at 2μm spacing intervals in the z-axis, and rendered in MetaMorph software (Molecular Devices). Images were further evaluated in ImageJ software (Schneider et al., 2012).

### Immunocytochemistry assay

Tow-cell embryos were fixed in methanol at −20°C overnight, washed in PBS (supplemented with 0.1% BSA), and incubated for 5 min in permeabilization solution (PBS, 0.1% BSA, 0.1% Triton X-100). Embryos were then incubated for 1 h in blocking solution (PBS, 0.1% BSA, 10% bovine fetal serum), following incubation with Monoclonal Anti-Cre antibody (Sigma-Aldrich Cat# C7988, RRID:AB_439697, 1:100 dilution in blocking solution) for 2 h at room temperature. After three PBS (0.1% BSA) washes, indirect detection was possible by immersing the embryos in blocking solution with Goat Anti-Mouse IgG1 (y1) Antibody, Alexa Fluor 594 Conjugated (Innovative Research Cat# 322588, RRID:AB_1502303, 1:500 dilution). Following three washes with PBS (0.1% BSA), embryos were immersed in blocking solution containing Mouse anti-Tubulin-Alexa-488 (Innovative Research Cat# 322588, RRID:AB_1502303, 1:25dilution) for 1 h. Embryos were washed in PBS (0.1% BSA) supplemented with NucBlue Fixed Cell Stain ReadyProbes (Invitrogen), and preserved with Prolong Gold (Invitrogen). Embryos were scanned with a Fluoview FV1000 confocal microscope (Olympus), with the aid of a 40× oil objective.

### Assessment of color-coded distribution of cells in blastocysts

Blastocyst images were visually inspected by one investigator. Cells were identified with the by their nucleus staining with DAPI. The cells were classified according to two criteria: First, cells were assigned to the embryonic (em) or abembryonic (abe) pole of the embryo. Second, the cells were color coded as red, green or no-color.

### Single-cell RNA-sequencing

Four- (n=9) and 8-cell (n=4) embryos imaged and having two or more cells expressing at least one of the fluorescent proteins (RFP or GFP) were subjected to single-cell RNA-sequencing. Embryos were briefly treated with Tyrode’s acid solution (Sigma) for removal of zona pelucida. The embryos were washed in PBS and incubated in Trypsin solution (Invitrogen) with RNase inhibitor (1lU/μl, Clontech) and BSA (50μg/μl) for the separation of the blastomeres. The blastomeres were separated by gentle pipetting, washed in PBS supplemented with RNase inhibitor and BSA, and snap frozen in individual micro-centrifuge tubes.

We amplified the cDNA of individual blastomeres according to the SMART-seq2 protocol (Picelli et al., 2014), followed by library preparation using Nextera XT DNA Sample Prep Kit (Illumina). The yields of cDNA and library amplification were assessed by Qubit (Invitrogen), and the quality of the distribution was assessed by Bioanalyzer (Agilent). Libraries were sequenced to produce pair-end reads of 100 nucleotides each in a HiSeq 2500 sequencer (Illumina).

### Alignment of sequencing reads to fluorescent protein genes in brainbow construct v2.1

Sequencing reads from each cell were aligned to DNA sequences of the four fluorescent protein genes present on brainbow construct v2.1 (Addgene plasmid repository 18723) using STAR aligner. Reads that aligned exclusively to one of the sequences were used for counting. Cells were assigned a color coding (RFP, YFP, CFP or GFP) according to the greatest number of reads matching to one of the fluorescent proteins, or NA when no read aligned to any of the DNA sequences.

### Alignment to the reference genome

STAR (STAR_2.5.1b, default parameters) was used to align single-cell sequencing reads to the reference genome (mm10, UCSC). Only uniquely aligned reads were used for the following analyses. Annotation files for Refseq genes and RepeatMasker defined repeats were downloaded from UCSC (February 2015), which contained a total of 24,225 genes and 5,138,231 repeats, respectively. These annotation files and the uniquely mapped reads in each cell were given as input to HTSeq-count (version 0.9.1) for counting of reads associated with each gene or repeat.

### Principle component analysis

Principle component analysis was carried out based on all the genes that exhibited FPKM>1 in at least one blastomere.

### Analysis of variance (ANOVA) analysis

A generalized linear model was fit to every gene as follows:

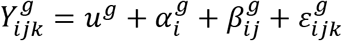

where 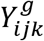 is the observed expression level (log2(FPKM+1)) of gene *g* in embryo *i*, lineage *j* (*j* = 0,1), cell *k*; *u*^*g*^ is the average expression level of gene *g* in all cells; 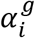 is the embryo-effect to average expression; 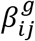 is the difference of expression levels between the two cell division lineages; 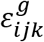 is the error term. R package {stats} was used to implement this linear model and test for every gene the null hypothesis that no reproducible expression differences between the two division, namely 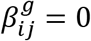. To account for multiple hypothesis testing, R package {qvalue} was used to compute q-values from ANOVA derived p-values.

ANOVA analysis for TRENIs was carried out in the same way as for genes, except that index *g* in the above generalized linear model was used as the index for TRENIs.

### Comparison of lineage variation and embryo variation

For every gene (*g*) we calculated the ratio (*r*^*g*^) of lineage sum of squares to combined sum of squares of embryo and lineage as

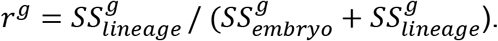

The random variable (*R*) of ratio of lineage sum of squares and combined sum of squares of embryo and lineage was defined as

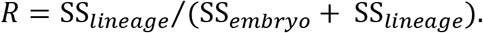

### Identification of novel transcript isoforms

Novel transcript isoforms were identified by Cufflinks (v2.2.1 with -g parameter) (Trapnell et al., 2012) with all Rainbow-seq data. Repeat sequences that overlapped with any Refseq annotated exons were removed from further analysis. TRENIs were identified by intersecting the remaining repeats with novel transcript isoforms using bedtools (2.27.0) (Quinlan and Hall, 2010).

### Calculation of junction ratio

For each repeat sequence (indexed by *i*), HTSeq-count (version 0.9.1) was used to count the number of reads mapped to this repeat (*T*_*i*_) and the number of reads spanning the junction of this repeat and nearby exons (*S*_*i*_). Junction ratio (*JR*) for the *i*^th^ repeat was calculated as
*JR*_*i*_ = *T*_*i*_/*S*_*i*_.

### Analysis of association between repeat families and TRENIs

Repeat families and the correspondence of individual repeat sequences to repeat families were retrieved from RepeatMasker (February 2015). A contingency table was built for each repeat family and TRENIs as follows.

|  | Contained within TRENIs | Not contained in TRENIs |
| --- | --- | --- |
| Is a member of this family | $a$ | $b$ |
| Not a member | $c$ | $d$ |

In the above table *a, b, c, d* are counts of repeat sequences and they sum to the total number repeat sequences (5,138,231). Odds ratio representing the degree of association between this repeat family and TRENIs was calculated as

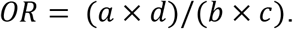

Ninety five percent confidence interval for odds ratio was calculated as

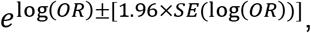

where 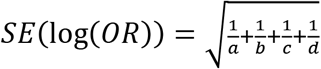

## SUPPLEMENTARY FIGURES

**Figure S1.**
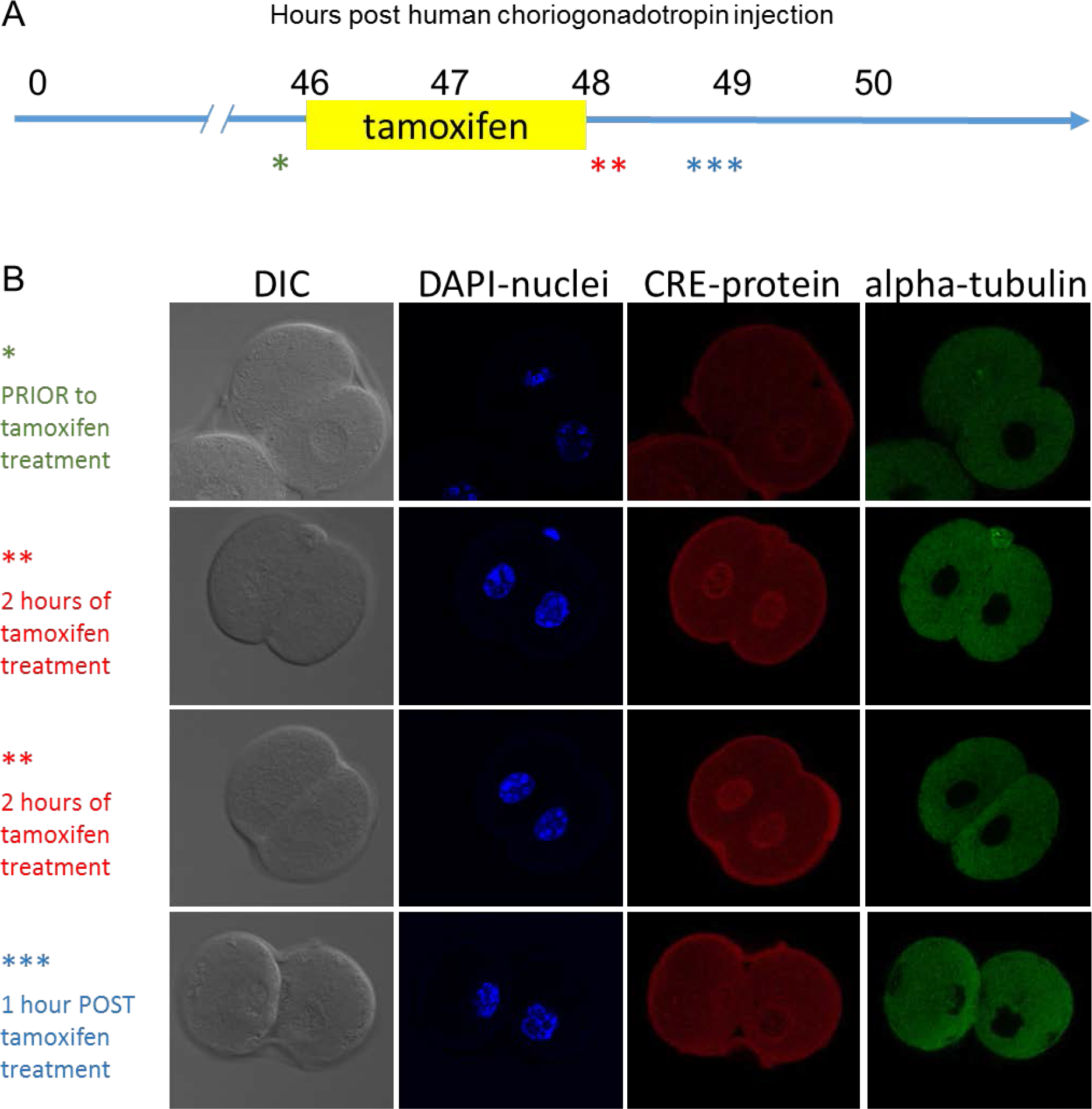
Tamoxifen induced nuclear localization of Cre recombinase.

**Figure S2.**
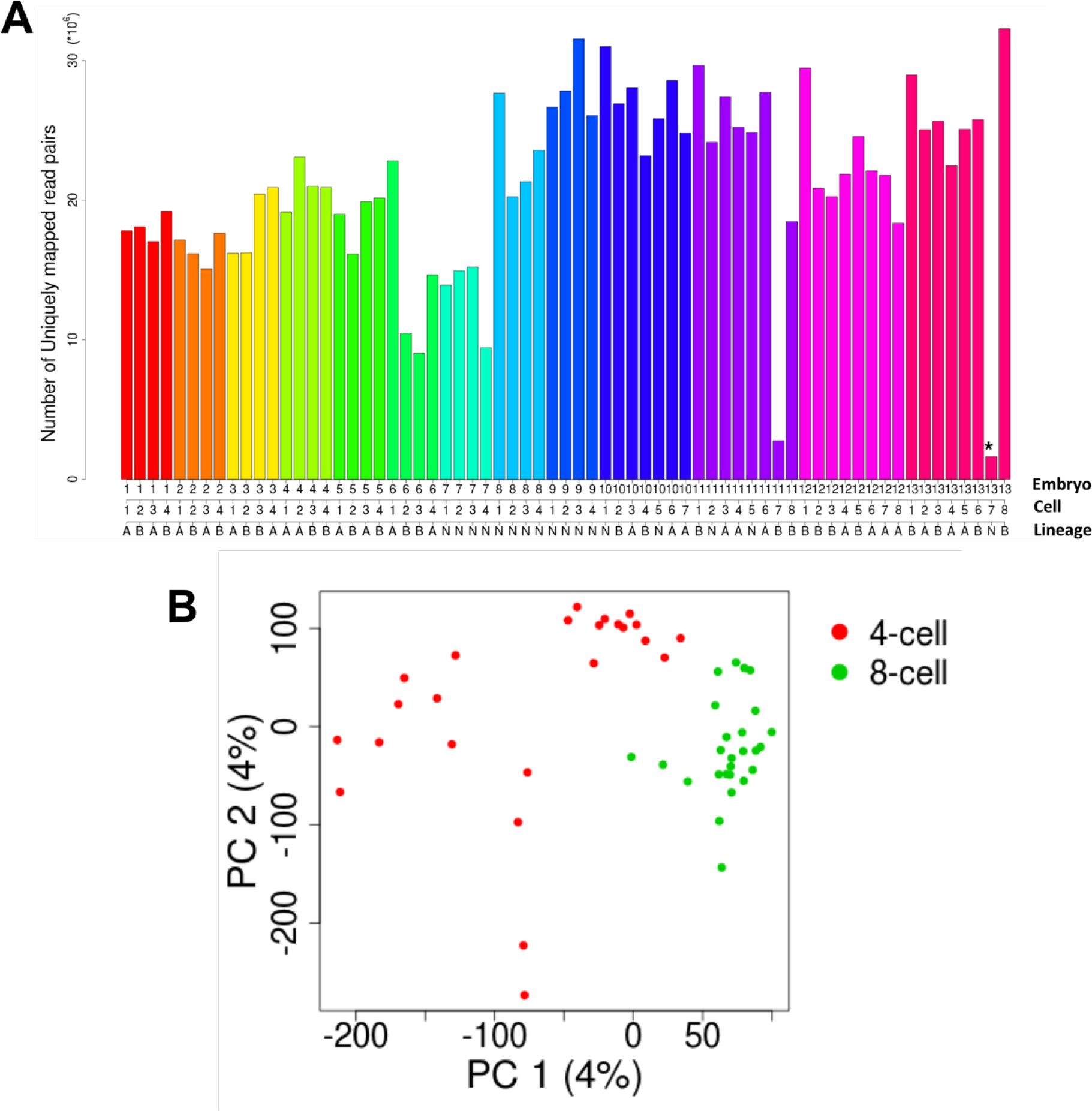
Overview of the samples. (A) Number of uniquely mapped read pairs (y axis) from each single cell (column). Each single cell is marked with an embryo number (Embryo), a cell number (Cell) and a binary lineage indicator (A or B). N: unresolved. (B) Principle component analysis of lineage resolved blastomeres at 4- (red) and 8-cell stages (green), using all genes with FPKM>1 in a least one blastomere.

**Figure S3.**
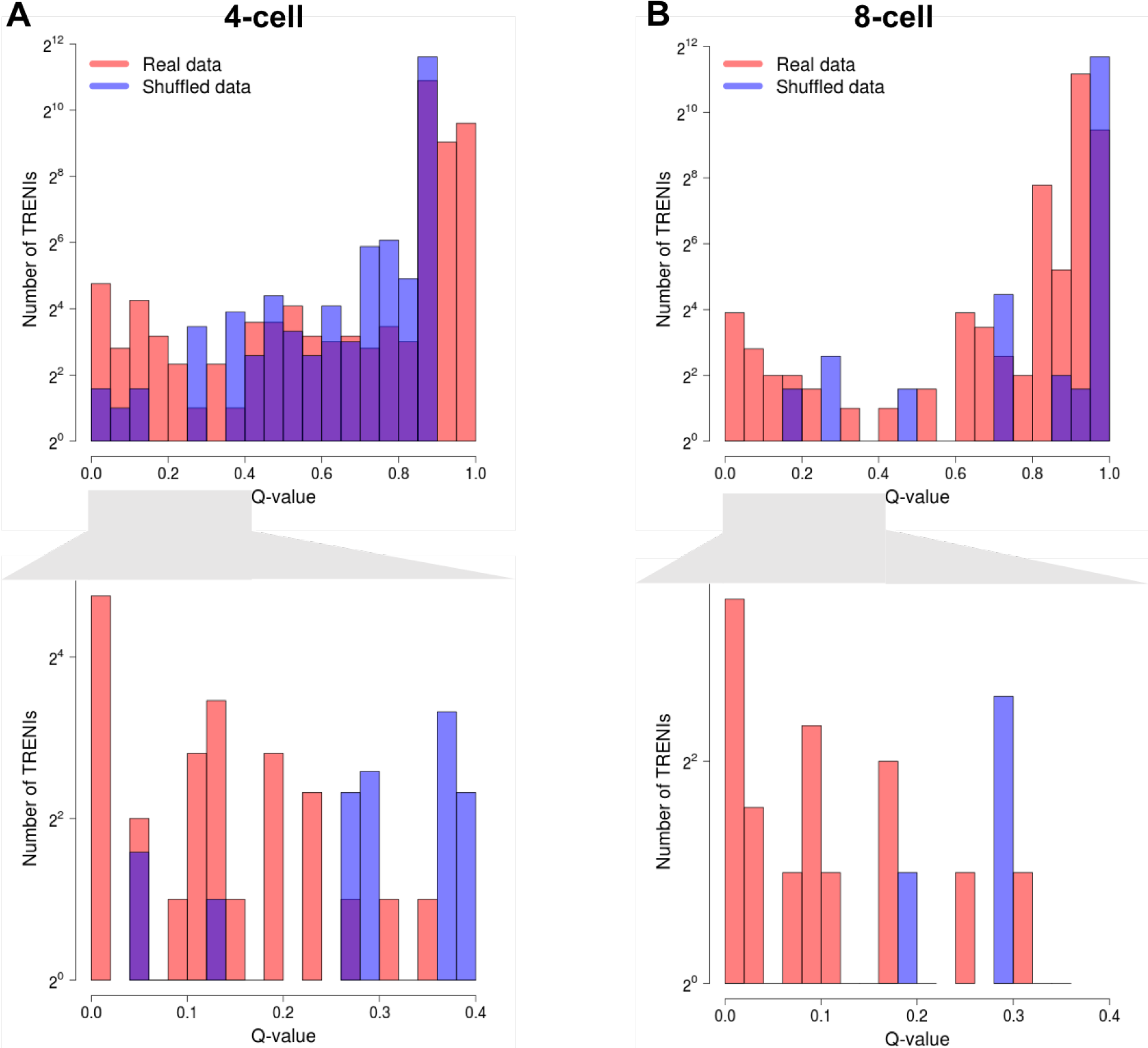
Histogram of TRENIs’ q-values derived from tests of equivalent expression between lineages at 4- (A) and 8-cell (B) stages, based from real data (red) and shuffled data (blue). Lower panels: expansion of the range 0 – 0.4 showing difference of real data and shuffled data in the range of small q-values.

**Figure S4.**
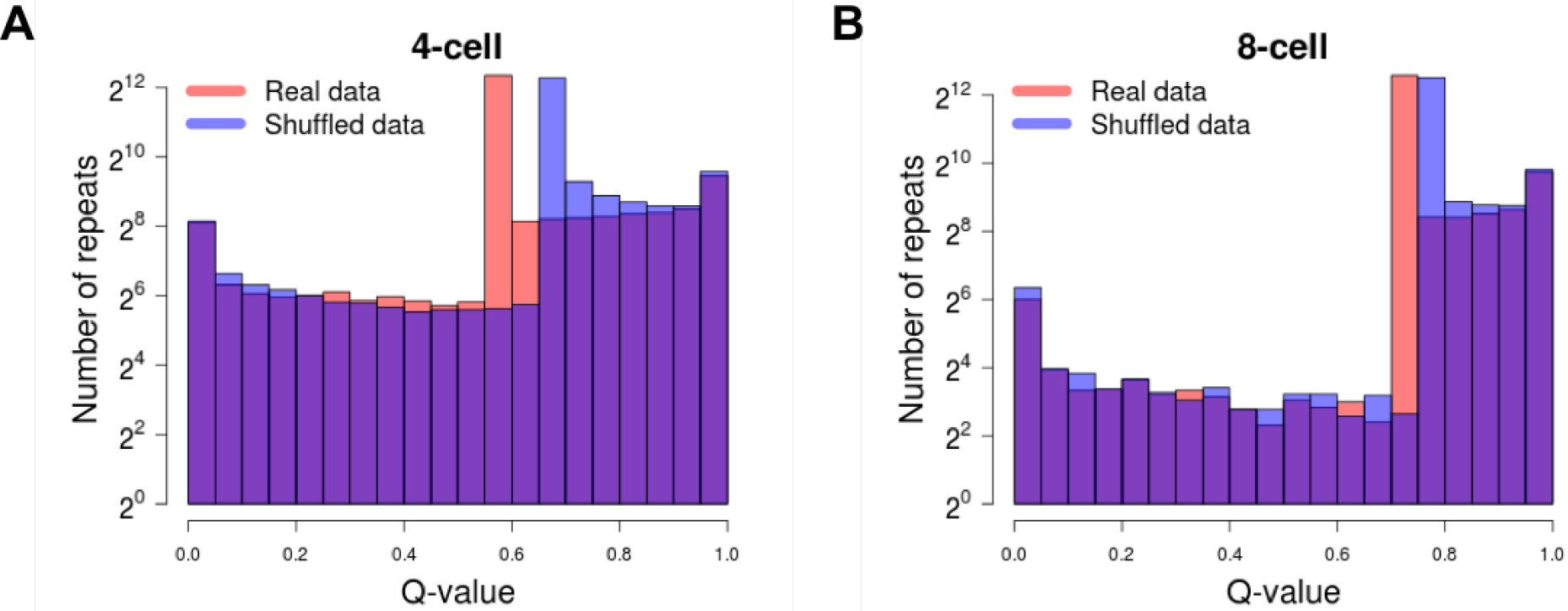
Tests of equivalent expression between lineages for all expressed repeat sequences. Repeats’ q-values derived from real (red) and shuffled data (blue) exhibited similar histograms, in both 4- (A) and 8-cell (B) stages.

**Figure S5.**
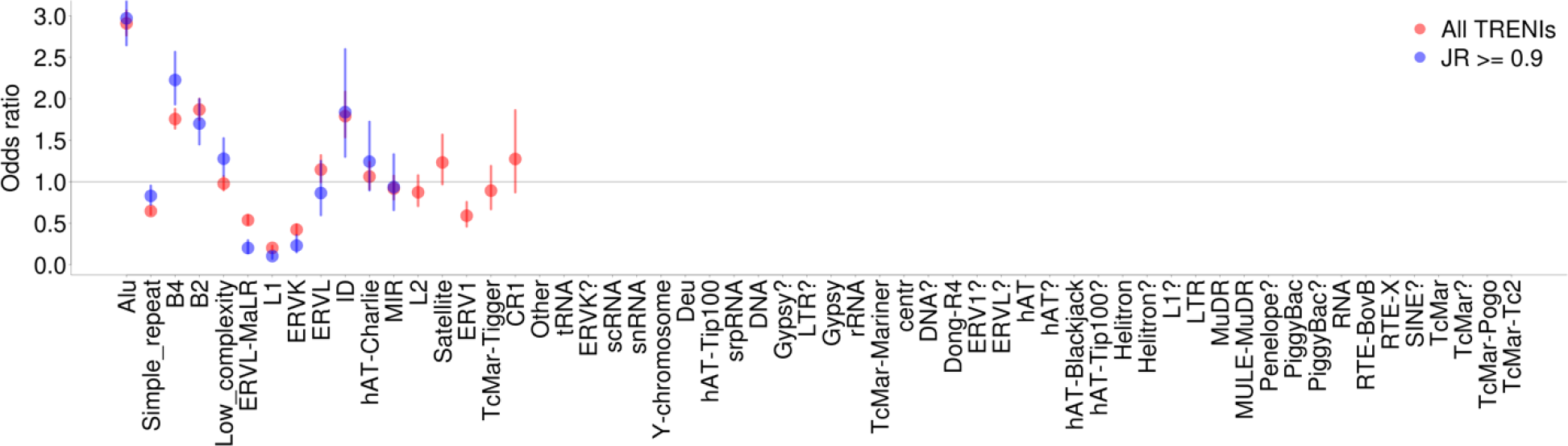
Sensitivity analysis of transposon related novel isoforms (TRENIs). Odds ratios between TRENIs and each transposon family (columns), calculated from only the repeat sequences with junction ratio (JR) > 0.9 (blue) were not very different from those calculated from all TRENI embedded repeats (red). Odds ratio was not calculated when the TRENIs of a repeat family were fewer than 20. Vertical bars: 95% confidence intervals.

**Figure S6.**
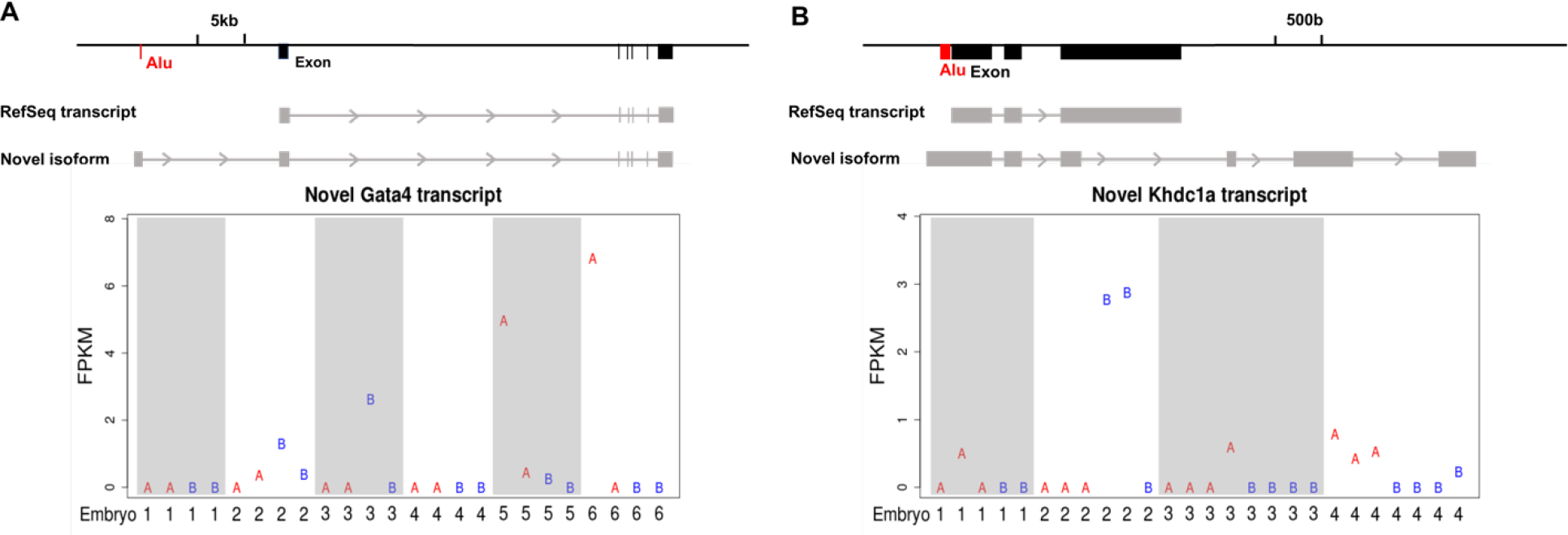
Novel transcript isoforms of *Gata4* and *Khdc1a* containing murine Alu in 5’ UTRs. (A) Expression levels of a novel transcript isoform of *Gata4* in 4-cell blastomeres. (B) A novel transcript isoform of *Khdcla* in 8-cell blastomeres. FPKM (y axis) of every blastomere (column) in each embryo (marked by embryo number in columns). Shaded columns delineate different embryos. The blastomeres of the two lineages are marked with A (red) and B (blue), respectively.

## SUPPLEMENTARY TABLES

**Table S1.**
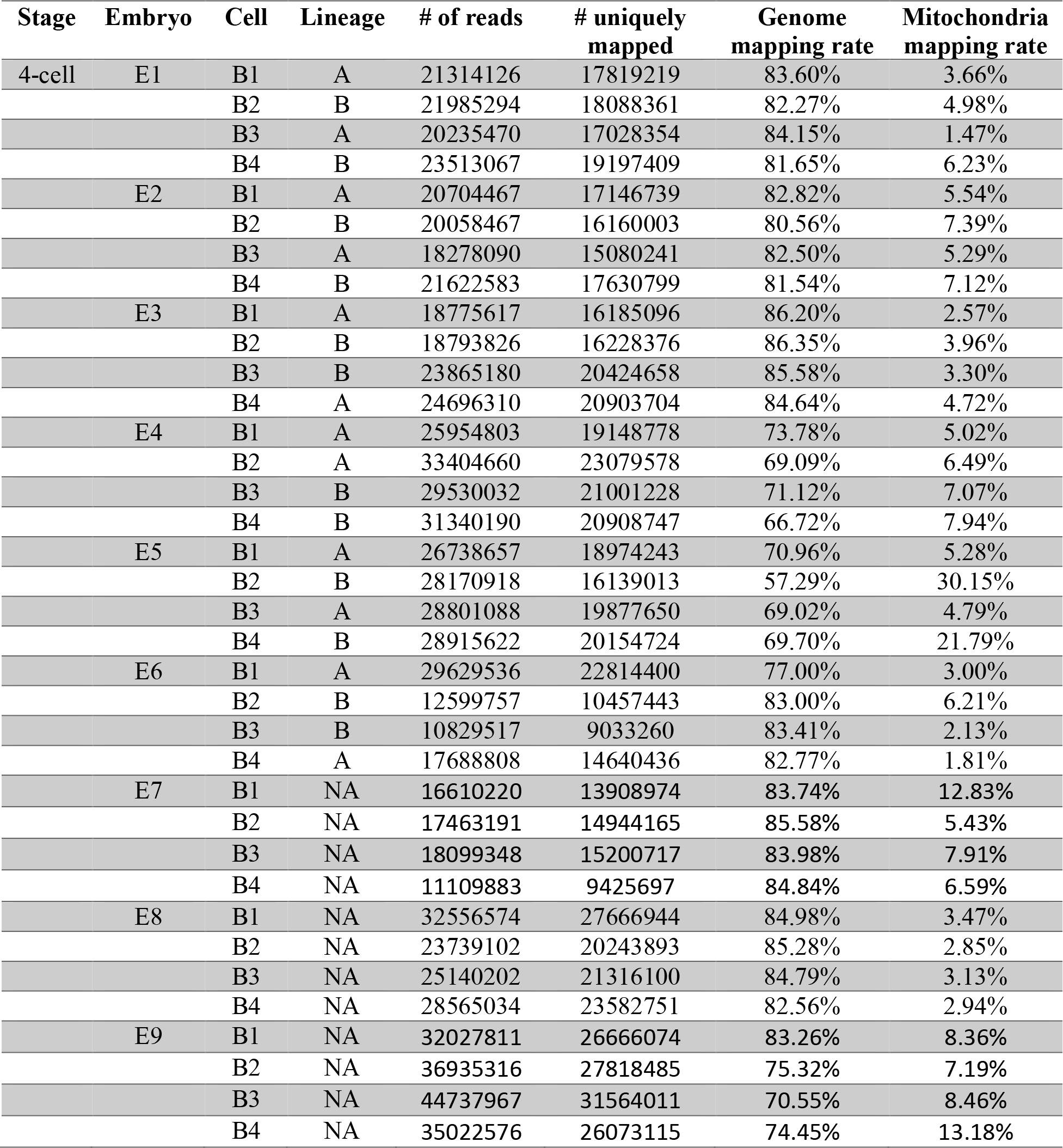
Summary of Rainbow-seq data from 4-cell stage blastomeres. Each blastomere (row) is indexed by embryonic stage (Stage), embryo number (Embryo), cell number (Cell), cell division lineage (Lineage), the total number of Rainbow-seq reads (# of reads), number (# uniquely mapped) and the percentage (Genome mapping rate) of reads uniquely mapped to the mm10 genome, and the percentage of reads mapped to the mitochondra genome (Mitochondria mapping rate). NA: undetermined.

| Stage | Embryo | Cell | Lineage | # of reads | # uniquely mapped | Genome mapping rate | Mitochondria mapping rate |
| --- | --- | --- | --- | --- | --- | --- | --- |
| 4-cell | E1 | B1 | A | 21314126 | 17819219 | 83.60% | 3.66% |
|  |  | B2 | B | 21985294 | 18088361 | 82.27% | 4.98% |
|  |  | B3 | A | 20235470 | 17028354 | 84.15% | 1.47% |
|  |  | B4 | B | 23513067 | 19197409 | 81.65% | 6.23% |
|  | E2 | B1 | A | 20704467 | 17146739 | 82.82% | 5.54% |
|  |  | B2 | B | 20058467 | 16160003 | 80.56% | 7.39% |
|  |  | B3 | A | 18278090 | 15080241 | 82.50% | 5.29% |
|  |  | B4 | B | 21622583 | 17630799 | 81.54% | 7.12% |
|  | E3 | B1 | A | 18775617 | 16185096 | 86.20% | 2.57% |
|  |  | B2 | B | 18793826 | 16228376 | 86.35% | 3.96% |
|  |  | B3 | B | 23865180 | 20424658 | 85.58% | 3.30% |
|  |  | B4 | A | 24696310 | 20903704 | 84.64% | 4.72% |
|  | E4 | B1 | A | 25954803 | 19148778 | 73.78% | 5.02% |
|  |  | B2 | A | 33404660 | 23079578 | 69.09% | 6.49% |
|  |  | B3 | B | 29530032 | 21001228 | 71.12% | 7.07% |
|  |  | B4 | B | 31340190 | 20908747 | 66.72% | 7.94% |
|  | E5 | B1 | A | 26738657 | 18974243 | 70.96% | 5.28% |
|  |  | B2 | B | 28170918 | 16139013 | 57.29% | 30.15% |
|  |  | B3 | A | 28801088 | 19877650 | 69.02% | 4.79% |
|  |  | B4 | B | 28915622 | 20154724 | 69.70% | 21.79% |
|  | E6 | B1 | A | 29629536 | 22814400 | 77.00% | 3.00% |
|  |  | B2 | B | 12599757 | 10457443 | 83.00% | 6.21% |
|  |  | B3 | B | 10829517 | 9033260 | 83.41% | 2.13% |
|  |  | B4 | A | 17688808 | 14640436 | 82.77% | 1.81% |
|  | E7 | B1 | NA | 16610220 | 13908974 | 83.74% | 12.83% |
|  |  | B2 | NA | 17463191 | 14944165 | 85.58% | 5.43% |
|  |  | B3 | NA | 18099348 | 15200717 | 83.98% | 7.91% |
|  |  | B4 | NA | 11109883 | 9425697 | 84.84% | 6.59% |
|  | E8 | B1 | NA | 32556574 | 27666944 | 84.98% | 3.47% |
|  |  | B2 | NA | 23739102 | 20243893 | 85.28% | 2.85% |
|  |  | B3 | NA | 25140202 | 21316100 | 84.79% | 3.13% |
|  |  | B4 | NA | 28565034 | 23582751 | 82.56% | 2.94% |
|  | E9 | B1 | NA | 32027811 | 26666074 | 83.26% | 8.36% |
|  |  | B2 | NA | 36935316 | 27818485 | 75.32% | 7.19% |
|  |  | B3 | NA | 44737967 | 31564011 | 70.55% | 8.46% |
|  |  | B4 | NA | 35022576 | 26073115 | 74.45% | 13.18% |

**Table S2.**
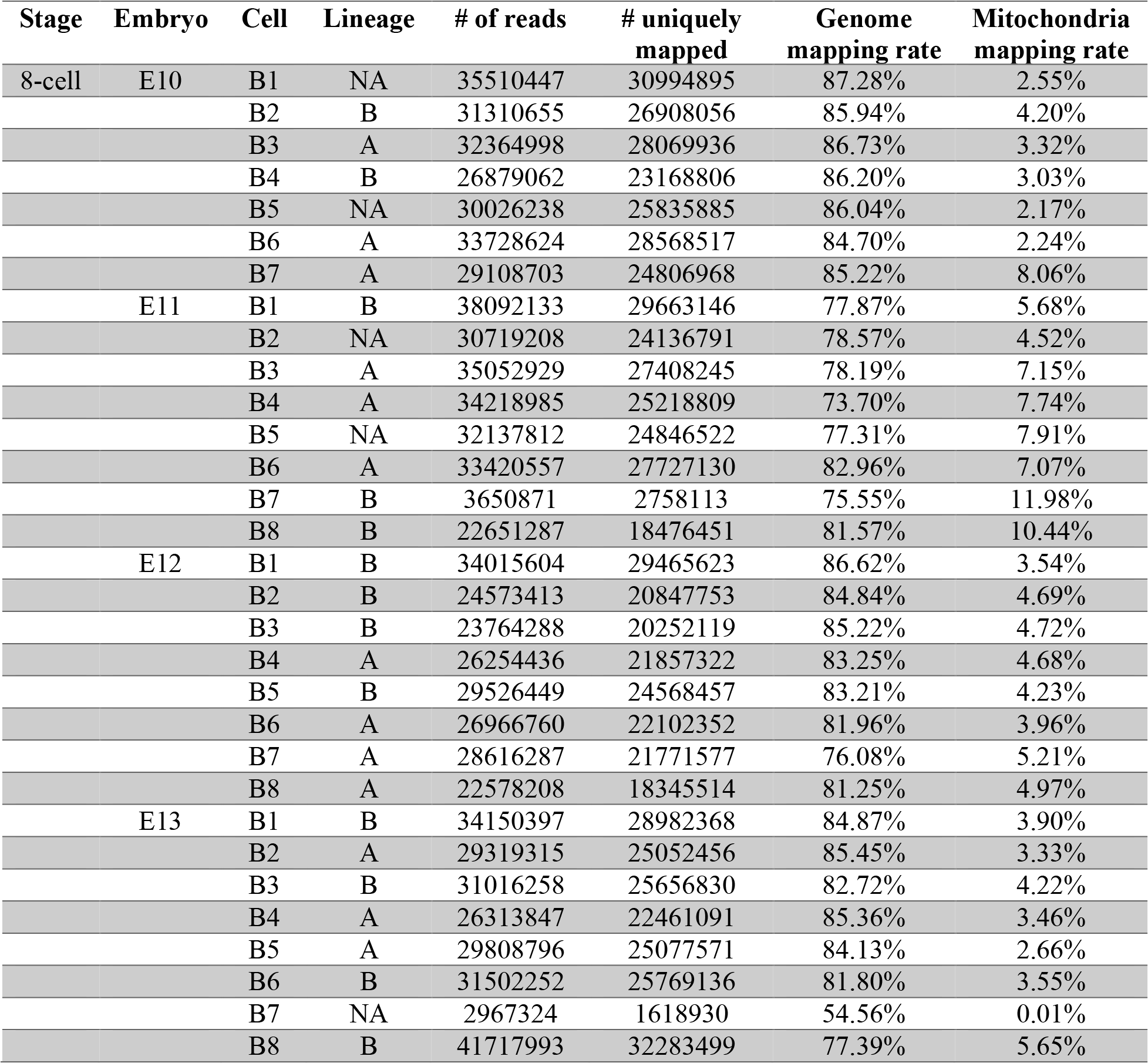
Summary of Rainbow-seq data from 4-cell stage blastomeres. Each blastomere (row) is indexed by embryonic stage (Stage), embryo number (Embryo), cell number (Cell), cell division lineage (Lineage), the total number of Rainbow-seq reads (# of reads), number (# uniquely mapped) and the percentage (Genome mapping rate) of reads uniquely mapped to the mm 10 genome, and the percentage of reads mapped to the mitochondra genome (Mitochondria mapping rate). NA: undetermined.

